# Ablation of the FACIT collagen XII disturbs musculoskeletal ECM organization and causes patella dislocation and myopathy

**DOI:** 10.1101/2021.12.29.474475

**Authors:** Mengjie Zhu, Fabian Metzen, Janina Betz, Mark Hopkinson, Juliane Heilig, Thomas Imhof, Anja Niehoff, David E. Birk, Yayoi Izu, Andrew A. Pitsillides, Janine Altmüller, Gudrun Schreiber, Mats Paulsson, Manuel Koch, Bent Brachvogel

## Abstract

Collagen XII, belonging to the fibril-associated collagens with interrupted triple helix (FACIT) family, assembles from three identical α-chains encoded by the COL12A1 gene. The trimeric molecule consists of three N-terminal noncollagenous NC3 domains joined by disulfide bonds followed by a short interrupted collagen triple helix at the C-terminus. Collagen XII is expressed widely in the musculoskeletal system and mutations in the COL12A1 gene cause an Ehlers-Danlos/myopathy overlap syndrome, which is associated with skeletal abnormalities and muscle weakness. Our study defines the role of collagen XII in patella development using the *Col12a1*^-/-^ mouse model. Deficiency in *Col12a1* expression causes malformed facies patellaris femoris grooves at an early stage, which leads to patella subluxation and growth retardation. Due to the patella subluxation, more muscle fibers with centralized nuclei occur in the quadriceps than in the gastrocnemius muscles indicating a local effect. To further understand the role of collagen XII in the skeletal tissues single cell RNAseq (scRNA-seq) was performed. Comparison of the gene expression in the tenocyte cell sub-population of wild type and *Col12a1*^-/-^ mice showed that several matrix genes are altered. Finally, we reinvestigated collagen XII deficient patients and observed a patella instability.

## Introduction

Extracellular matrix and, in particular, collagens play an important role in the musculoskeletal system. Collagen XII, a homotrimer which belongs to the subfamily of fibril-associated collagens with interrupted triple helices (FACIT), is widely expressed in mice and humans, e.g. in skin, cartilage, nerve, and muscle (1). In most tissues collagen XII is associated with collagen fibrils but also with tenascin-X (2, 3). Collagen XII occurs in several splice variants of which the largest is a 300 kDa protein (18 FN3, 4 VWA, 1 TSPN, and 2 COL domains), whereas the small splice variant lacks a large part of the N-terminal region (4). The larger form contains an additional heparin binding site in the 7th FN3 domain; moreover glycosaminoglycan chains are covalently attached. Collagen XII has been identified as a key organizer of the extracellular matrix in tendon, bone, cornea, and muscle (1). Mutations in the COL12A1 gene were identified as the cause of myopathic Ehlers-Danlos (mEDS) overlap syndrome (5, 6). Depending on the nature of the mutation, dominant or recessive, the phenotype varies in severity (7). Patients with a homozygous loss of function mutation show joint hyperlaxity combined with weakness precluding independent walking and feeding as newborn (8). Due to night-time hypoventilation, noninvasive night-time ventilation has been used. Severe kyphoscoliosis is another hall mark of mEDS. Interestingly, the muscle weakness is not progressive and patients with *de novo* missense mutations are more mildly affected. Surprisingly, the muscle weakness improves over time, including the ability to walk (5, 6).

*Col12a1*^*-/-*^ mice recapitulate the phenotype observed in COL12A1 deficient patients (6). However, as often observed for extracellular matrix (ECM) genes, the phenotypes in mice are less severe and, in the case of *Col12a1*^*-/-*^, the pups can feed by themselves and are able to walk. Analysis of the *Col12a1*^*-/-*^ bones revealed a brittle bone phenotype due to a lack of cell polarity of the osteoblasts. In addition to improperly formed bones, osteocytes have fewer dendritic extensions. This was observed by immunofluoresence microscopy and confirmed by transmission electron microscopy (9). Interestingly, these cell extensions are also drastically reduced in tenocytes of *Col12a1*^*-/-*^ mice, which causes a lack of clear fiber domain organization. Furthermore, abnormal fiber packing in *Col12a1*^*-/-*^ mice was observed by electron microscopy and causes a change in the biomechanics (9). Flexor digitorum longus tendons have a significantly larger cross-sectional area in *Col12a1*^*-/-*^ mice and exhibit increased stiffness. Detailed morphological analysis of different tendons revealed that 20– 60% of the *Col12a1*^*-/-*^ mice display discontinuity in the anterior, but not in the posterior cruciate ligament. Hence the absence of collagen XII increases the risk of anterior cruciate ligament injury (10).

The patella, which is superficially embedded within tendons, belongs to the subgroup of sesamoid bones. The patella increases the distance between the quadriceps muscle and the tibia and thereby the moment arm of the muscle, which enhances its extension force by up to 50% (11). In addition, the patella is essential for knee stability. Congenital pathologies, such as patella aplasia and hypoplasia, as well as surgical removal of the patella, reduce the ability to walk and cause osteoarthritis (12). Developmental studies revealed that the patella initially develops as part of the femur, but from a distinct pool of Sox9-and Scx-positive progenitor cells which are dependent on TGFβ - BMP signaling (13). Interestingly, mechanical load is required for the formation of the patellofemoral joint. In humans several genetic disorders affecting the patella have been identified. The first mutation of this class was in the TBX4 (T-box protein 4) gene and was found to cause ischiopatellar dysplasia, due to the essential role TBX4 as a transcription factor in lower limb development (14). Most of the genes causing patella malformation are linked to three major developmental processes: limb specification and joint pattern formation, DNA replication and chromatin, or bone development and regulation (15). In most cases the patella alteration goes along with multiple other congentital anomalies, e.g. ischiocoxopodopatellar syndrome, a rare autosomal dominant disorder characterized by a hypoplasia of the patella and various anomalies of the pelvis and feet (16).

Here, we address the physiological roles of collagen XII in the musculoskeletal system. First, we analyze the tissue distribution of collagen XII in the knee. Secondly, we determine the function of collagen XII in the muscle-tendon-knee unit of *Col12a1*^*-/-*^ mice and, by single cell RNAseq, define the molecular changes occurring in tenocytes upon loss of collagen XII. Finally we analyze the knees of COL12A1 deficient patients.

## Results

### Expression of collagen XII in the murine musculoskeletal system at different developmental stages

The musculoskeletal system, consisting of muscle, bone, cartilage, tendons, and ligaments, provides the physical support for weight and movements. Localization of collagen XII was analyzed by immunofluorescent staining of whole leg sections from infant (postnatal 3 days, 7 days) and juvenile (1 month) mice. This showed a wide distribution of collagen XII in the extracellular matrix of those tissues. Collagen XII was found in muscle, patella, quadriceps tendon and periosteum of the femur (Figure 1A). In the synovial joints, collagen XII was localized in the anterior and posterior cruciate ligaments and in the articular cartilage of femur and tibia (Figure 1B). Collagen XII was also detected in the enthesis, where the patella tendon attaches to the tibial plateau, as well as in the periosteum of the tibia (Figure 1C).

**Figure 1.**
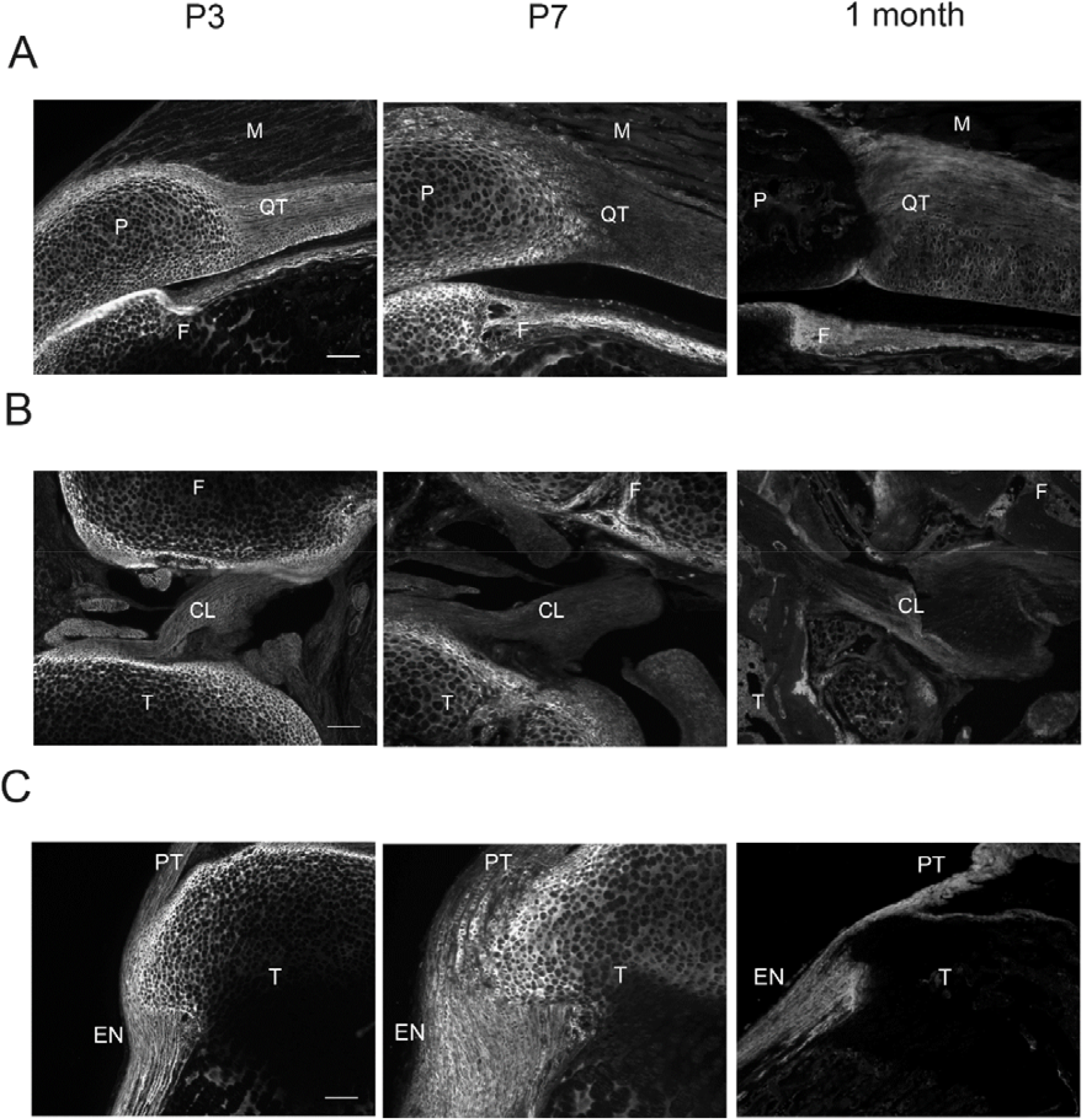
Expression of collagen XII during murine musculoskeletal system development. A) Localization of collagen XII was analysed by immunofluorescent staining in the whole leg of P3, P7 and 1 month old animals. Collagen XII has a wide distribution in patella (P), patella tendon (PT), femur (F), tibia (T) cartilage, cruciate ligament (CL), quadriceps tendon (QT), enthesis (EN), and muscle (M). B) Close up of the collagen XII expression in patella, cruciate ligament and enthesis. Scale bar: 100 μm.

### Patella subluxation in Col12a1^-/-^ mice

To determine the functions of collagen XII in the musculoskeletal system a *Col12a1*^*-/-*^ mouse model was used. This model was generated by targeted deletion of exon 2-5, as described previously (Izu, et al., 2011). Compared to their wild type littermate controls, KO mice showed growth retardation, as a decreased body length and weight was observed throughout development (Suppl. Figure 1A-B). A delay in secondary ossification center formation was found at P7 and P12, which relates to their shorter bones (Suppl. Figure 1C-F). Abnormal appearance of the stifle joint was detected at 3 months (Figure 2A). μCT analysis was performed on the distal hind limb joints of 1 month old *Col12a1*^*-/-*^ mice and indicated subluxation and growth retardation of the patella (Figure 2B). Histological analysis with H&E and Alcian Blue staining conducted in serial section planes, showed that at P3 the sections were comparable between genotypes and patella subluxation was not detected in most KO mice. At P12, at the level of femoral secondary ossification center formation, the patella fitted in the facies patellaris femoris groove in wild type animals, while in the KO animals the patella could not be found at the proper location. This was similar in 1 month old mice, where in WT mice a well ossified patella was found on top of the femur, while in KO mice the patella remained cartilaginous and was further removed (Figure 2C). At P3 only 20% of KO animals (2 out of 10) displayed this subluxation phenotype, while at P12 and 1 month of age, the incidence rose to 75%, suggesting that the subluxation increases gradually during development with increased mobility (Figure 2C-D).

**Figure 2.**
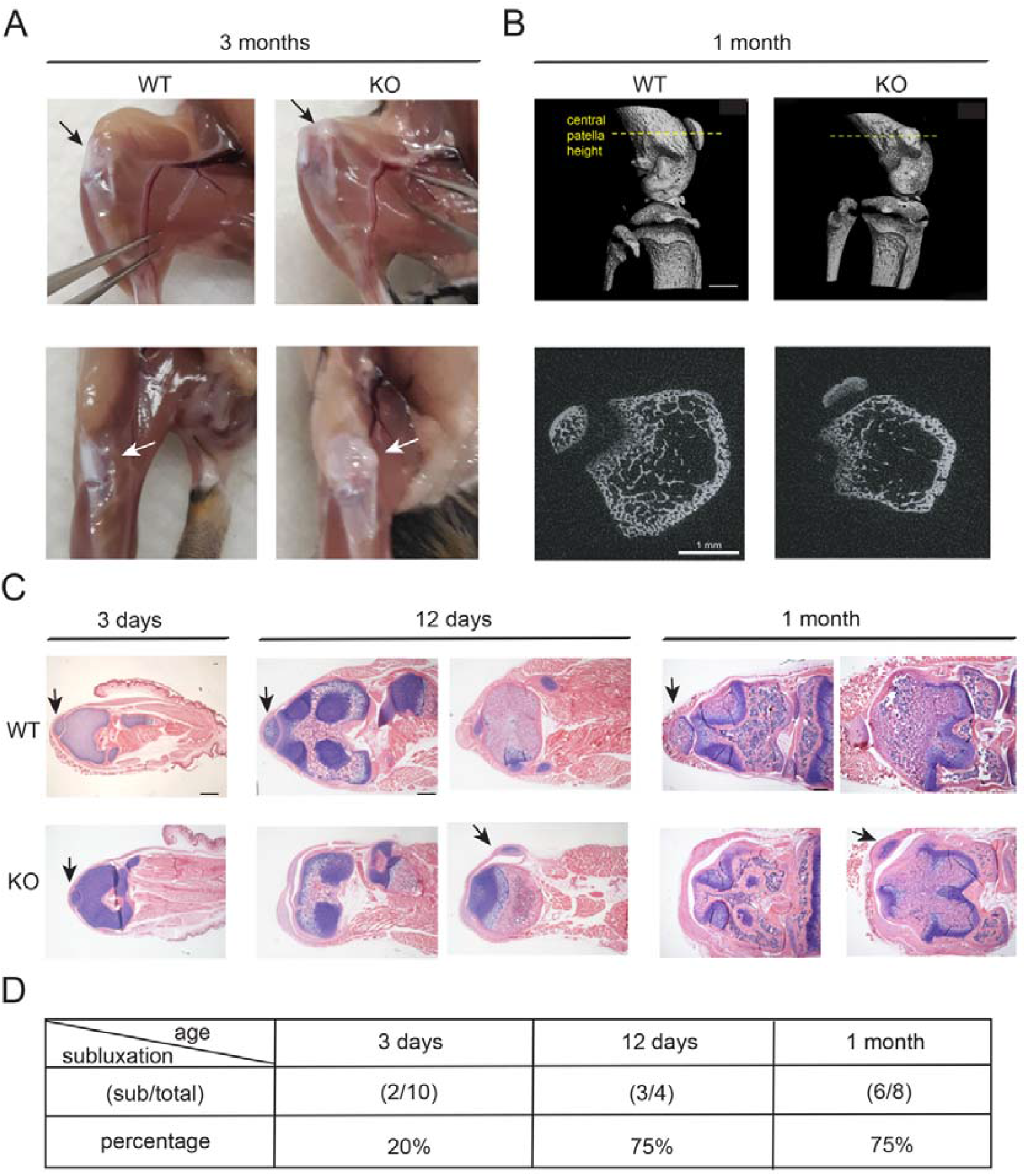
Patella subluxation in *Col12a1*^*-/-*^ mice. A) Representative pictures of anterior-posterior and subcutaneous views of 3 month old WT and *Col12a1*^*-/-*^ mouse joints. B) μCT analysis of 1 month old WT and KO mouse joints (n=4). Scale bar: 1 mm. C) In plane matched paraffin sections from WT and *Col12a1*^*-/-*^ mice were stained with hematoxylin (nuclei, purple), eosin (cytoplasm, pink) and alcian blue (proteoglycans, blue). Arrows indicate the location of the patella. Scale bar: 500 μm. D) Patella subluxation rate in WT and *Col12a1*^*-/-*^ mice of different ages.

### Col12a1^-/-^ mice display quadriceps muscle alteration

A profound reduction in quadriceps muscle mass was found in *Col12a1*^*-/-*^ mice compared to WT littermates at P12, 1 month and 6 months of age. (Figure 3A). This correlates with the occurrence of patella subluxation. To determine whether the loss of muscle mass is specific for the quadriceps and due to patella subluxation, the gastrocnemius muscle, as one of the tibial muscles, was used as a control. At 6 months of age, both quadriceps and gastrocnemius muscle mass was significant decreased in KOs when normalized to the body weight of the individual mouse (Figure 3B). In *Col12a1*^*-/-*^ mice, an increased number of muscle fibers with centralized nuclei and a larger fiber diameter was found in the quadriceps (Figure 3D, E), when H&E staining was performed on cross-sections of quadriceps and gastrocnemius muscles at 6 months of age (Figure 3C). In the gastrocnemius, there was no significant difference in the number of muscle fibers with centralized nuclei, only a slight increase in muscle fiber diameter. This indicates that the major muscle phenotype is specific for the quadriceps muscle.

**Figure 3.**
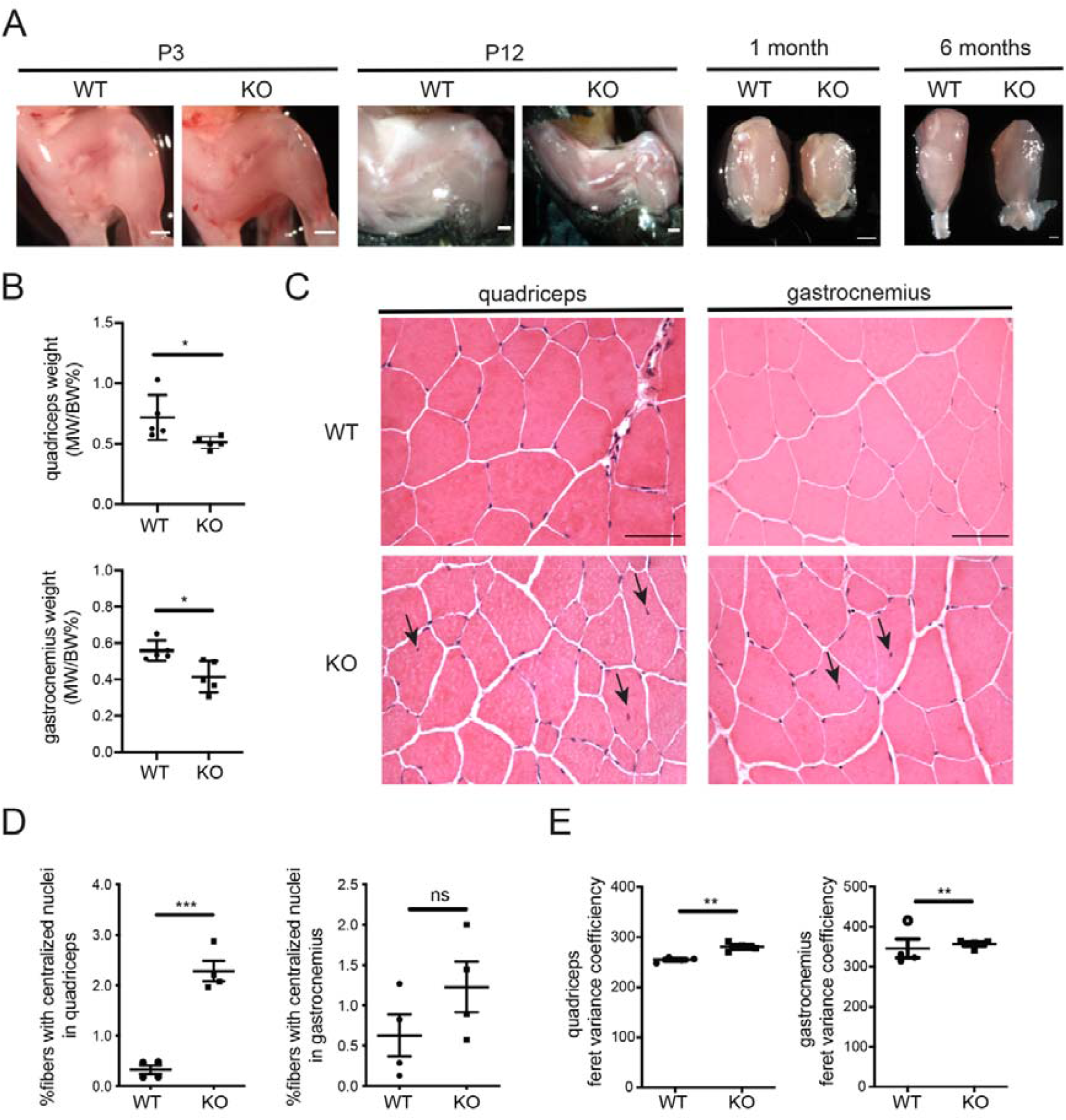
Col12a1^-/-^ mice show muscle alterations. A) Representative pictures of the hind limbs of P3 and P12 animals after stripping the skin and quadriceps muscle isolated from 1 month and 6 months mice. B) Quadriceps and gastrocnemius muscles were isolated from 6 months old mice and their weight normalized by individual body weight. MW: muscle weight, BW: body weight. (n=5). C-E) In plane matched 7 μm cryo cross sections of quadriceps and gastrocnemius muscle from WT and *Col12a1*^*-/-*^ (n=4) of 6 month old mice were stained with hematoxylin (nuclei, purple) and eosin (cytoplasm, pink). Scale bar, 100 μm. Black arrowheads indicate the centralized nuclei. Centralized nuclei number given as the percentage of total muscle fibre number in quadriceps and gastrocnemius muscles. Feret variance coefficient measurements determined with the minimal ‘Feret’s diameter’ of the muscle. Grubb’s test (alpha=0.05) was carried in all data sets to exclude outliers. One sample is excluded (labelled as white circle) in the gastrocnemius Feret variance coefficiency as an outlier. *, p < 0.05; **, p < 0.01, ***, p < 0.001. Error bars are mean ± SEM.

### Facies patellaris femoris groove malformation in Col12a1^-/-^ mice

Distal hind limbs from WT and KO mice at P1 were stained in Lugol’s solution for soft tissue visualization and quantification by X-ray based μCT scanning. The original data were reconstructed, showing that the femur (grey), the rectus femoris muscle (green), and the patella and the surrounding ligament (red; Figure 4B). Among the three mice in the *Col12a1*^*-/-*^ group a patella subluxation was found in two. Statistical analysis of the reconstructed rectus femoris showed a reduced length and volume in KO group compared to WT (Figure 4C). Volume renderings of μCT data revealed malformed facies patellaris femoris grooves in all KO mice (Figure 4A). In WT joints, facies patellaris femoris grooves were well-formed and appeared to have concave surfaces, while in all KO joints they had convex surfaces. KO mice also showed a trend towards a lower rectus femoris length and a perimeter at the mid-shaft of the femur which was significantly smaller than in WT mice (Figure 4C).

**Figure 4.**
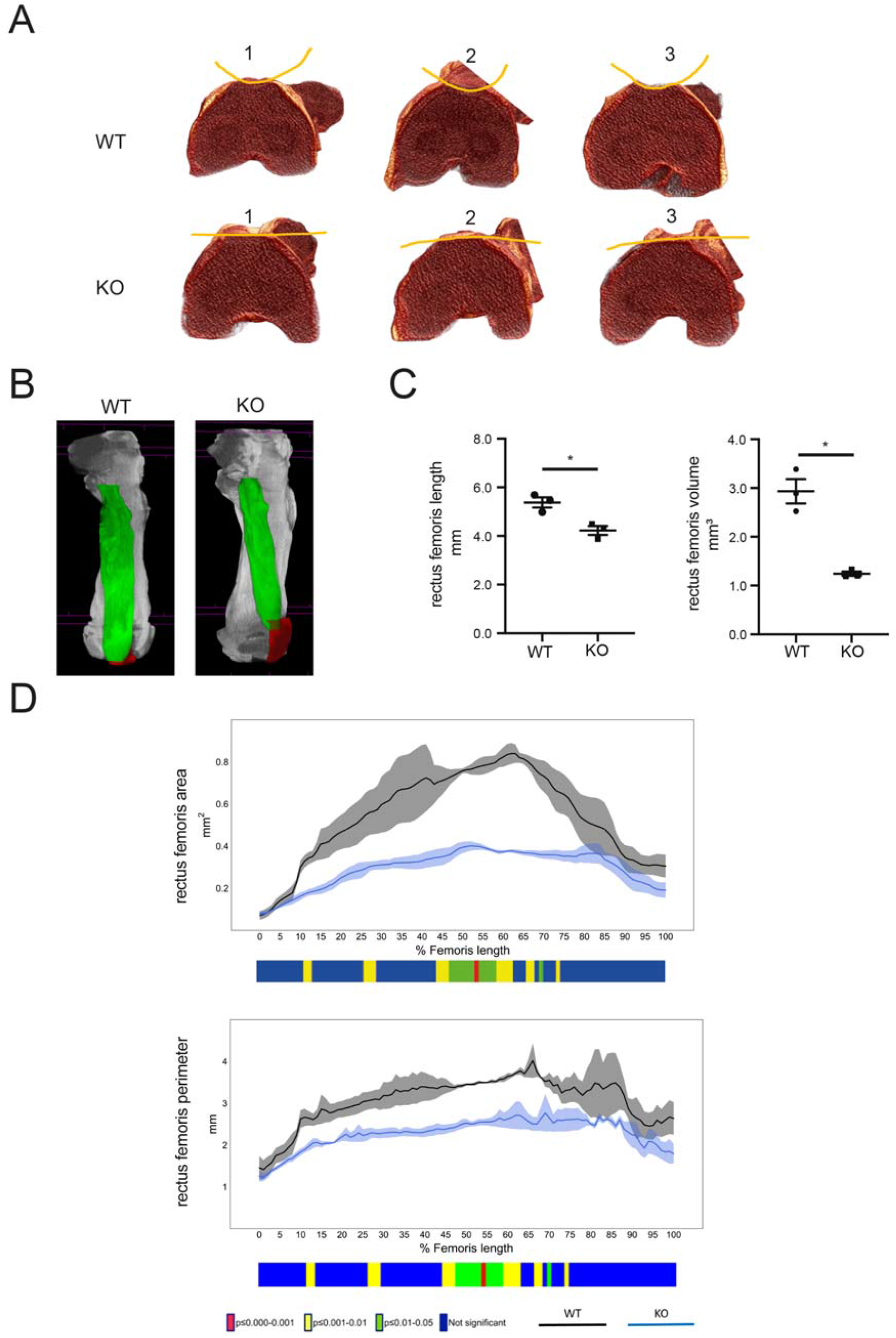
*Col12a1*^*-/-*^ mice show a malformed facies patellaris femoris groove and rectus femoris alterations at P7. A) Representative images of μCT volume renderings of femora from WT and KO mice (n=3) demonstrating the malformed facies patellaris femoris groove indicated by the yellow lines. B) Representative images of μCT volume renderings of femora (grey), rectus femoris (green), patella and ligament (red) from WT and KO mice at P7 (n=3). C) 3D morphometric analysis shows that KOs have reduced rectus femoris length and volume. Error bars are mean ± SEM. *, p < 0.05; **, p < 0.01. D) μCT 2D morphometric analysis along the length of the Lugol’s solution stained rectus femoris from 0% (proximal end) to 100% (distal end) show differences between the KO and the WT groups (n=3) in muscle area and muscle perimeter. Line graphs represent the means ±SEM. Two-sample t-test was used to compare means between KO and WT groups. The graphical heatmap displays statistical differences at specific matched locations along the length of the femur. Red p≤0.000-0.001, yellow p≤0.001-0.01, green p≤0.01-0.05 and blue p≥0.05.

### Single-cell RNA-seq analysis of legs from Col12a1^-/-^ mice

Single cell RNA sequencing was used to elucidate changes in the transcriptome in individual cell subpopulations from whole hind limbs of a *Col12a1*^*-/-*^ and a WT mouse at P7. Single cell suspensions were obtained by collagenase digestion of the hind limbs and passed through cell strainers. Library construction of each cell was performed by amplifying reverse-transcribed cDNA, followed by labelling with a cellular barcode. The libraries were subjected to massive parallel sequencing. In the case of the WT animal 5490 cells with 17,577 mean reads per cell (1783 median genes per cell) were obtained, whereas for the *Col12a1*^*-/-*^ mouse 4212 cells with 19,596 mean reads per cell (1676 median genes per cell) were achieved. In total, 142 differentially expressed genes were identified. Marker genes were used to annotate and analyse cluster-specific gene expression. The collagen XII expressing cell populations were analyzed and low expression levels were found in the *Col12a1*^*-/-*^ cells as expected. To annotate individual cell clusters, differential gene expression analysis was used to identify cell type–specific marker genes. Clusters of immune cells (*Adgre1+*, macrophages; *S100a9+*, neutrophils; *Cc9a+*, B cells) were detected (Figure 5A) and here the *Col12a1* expression level was rather low. Tenocyte (*Tnmd+*), myoblast (*Myod1+*), and chondrocyte (*Col2a1+*) populations, as the major cell types in the musculoskeletal system, were recognized in both genotypes. In WT mice a high *Col12a1* expression level was found in tenocytes and chondrocytes, while only few myoblasts showed expression (Figure 5B). Next, a GO enrichment for the differentially expressed genes in the tenocyte cell population was performed and a strong correlation to the GO terms ECM, response to stress, and tissue development detected (Suppl. Fig. 3A-D). Further, differentially regulated genes in the tenocyte subpopulations of *Col12a1*^*-/-*^ and WT mice were identified. In total, 11 out of 274 genes in the core matrisome and 22 out of 836 matrisome-associated genes were differentialy expressed (Figure 6). Most of the ECM genes were down regulated in the *Col12a1*^*-/-*^ cells, whereas only one, inhibin beta A chain (Inhba), was upregulated. Collagen XI, microfibril-associated protein 4 (MFAP4), spondin-2 (SPON2) and Wnt inhibitory factor 1 (WIF1), a secreted protein that binds to Wnt proteins and inhibits Wnt activity, were all down regulated (Figure 6B-C). Interestingly, many cytokines, growth factors or proteases were also differentially expressed, however only in few *Tnmd+* tenocytes (Suppl. Fig 4).

**Figure 5.**
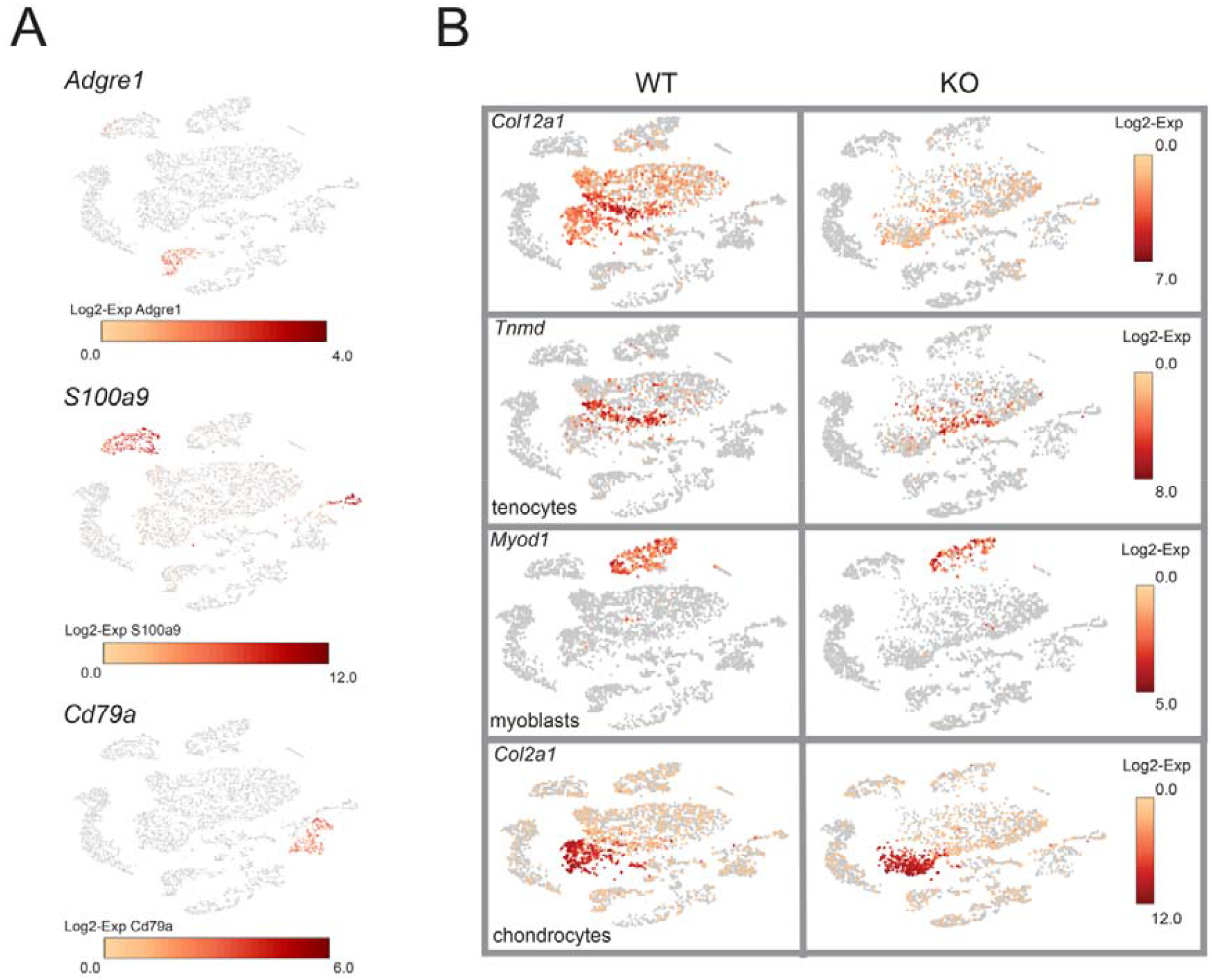
Identification of cell clusters based on marker genes. A) Expression of cell type specific marker genes was used to annotate individual cell clusters. Log2 expression intensity values of individual cells are indicated (color scale). *Adgre1* (F4/80), *S100a9, Cd79a* are used as markers for macrophages, neutrophils and B cells. B) *Col12a1*+, tenocyte (*Tnmd*+), myoblast (*Myod1*+), and chondrocyte (*Col2a1*+) cell clusters are shown in t-SNE plots.

**Figure 6.**
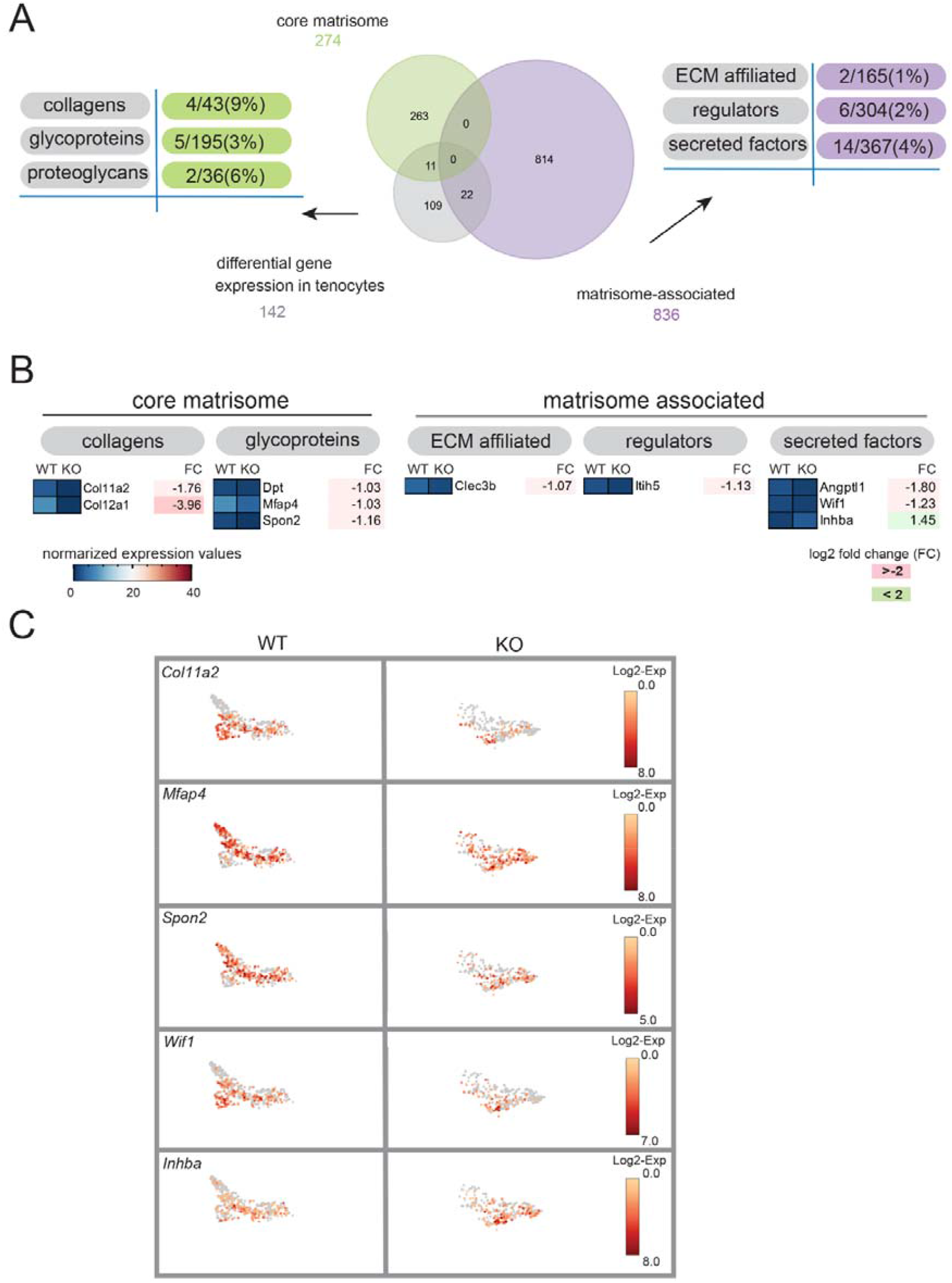
Matrisome analysis of the *Tnmd+* cell subpopulation at the transcriptomic level. A) The proportions of entities within the core matrisome or matrisome associated cluster are shown in a Venn diagram. The numbers and percentage of regulated genes found in subcategories are listed. B-C) List of selected genes based on fold change more than 1. Log2 intensity values and fold change between WT and *Col12a1*^-/-^ mouse are shown. Selected genes in *Tnmd+* subpopulation show different expression level in t-SNE plots

### Analysis of the human COL12A1 patella – knee unit

Homozygous recessive loss of COL12A1 in two siblings caused a severe myopathy accompanied with a hypermobility of joints. Both patients (now young adults) are not able to walk independently and require motorized wheelchairs. The myopathy did not progress since birth and remains stable. Since early on, both patients received night-time breathing support. They manage to change from a lying position to a sitting position, but never to stand up by themselves. Due to scoliosis, both patients as teenagers required back surgery for stabilization. In early childhood, very unstable knees were observed with very common patellar subluxations in both knees of both patients. X-ray analysis of their knees revealed in one knee a complete absence of the facies patellaris femoris groove, while the grooves were less pronounced in the others (Fig. 7). Not surprisingly, in the knee with the missing groove the patella cannot be kept in the proper position and, hence, was located on the side of knee. In addition, the patella of the left knee of the patient I is not well centred (Fig. 7A upper panel).

**Figure 7.**
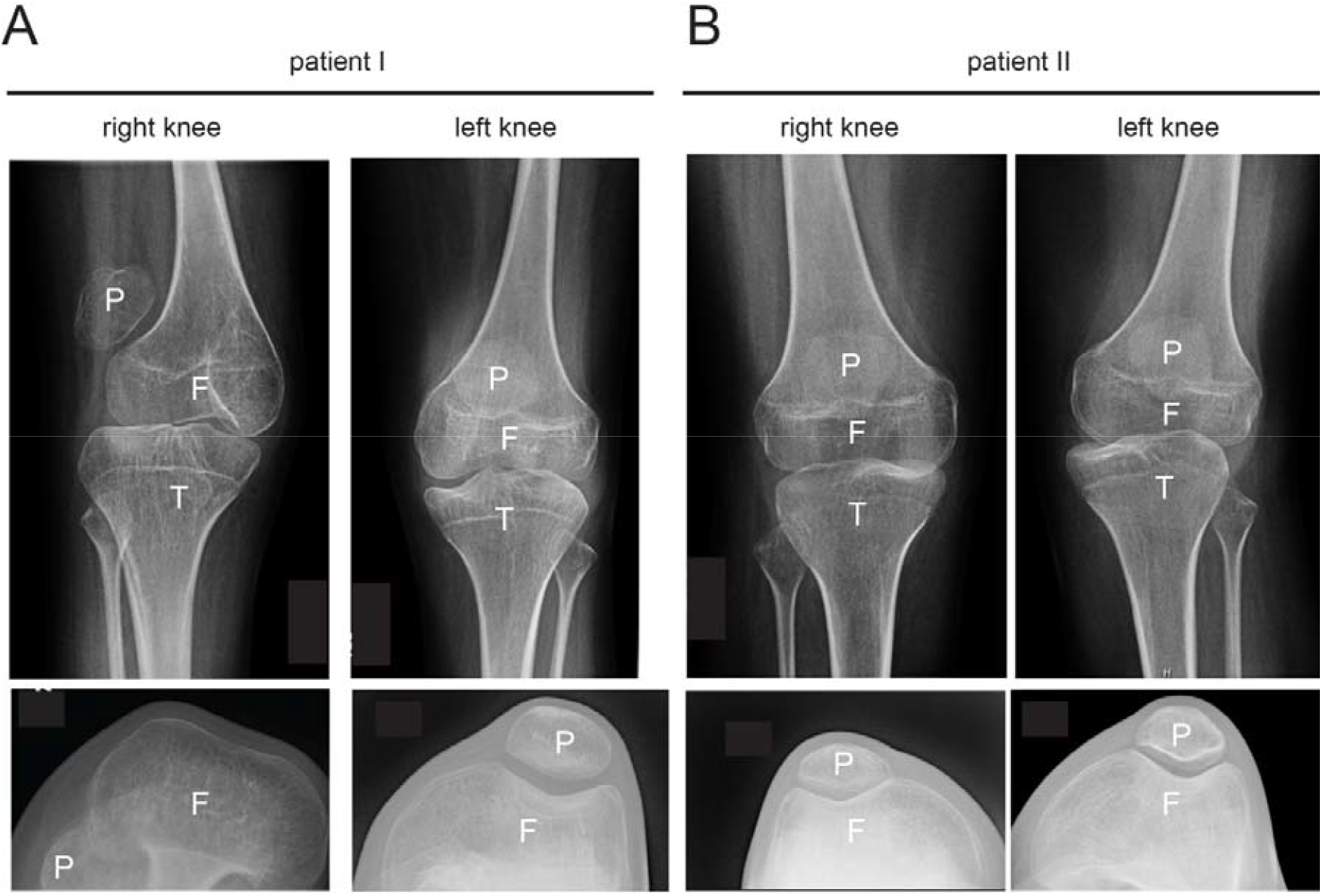
X-ray analysis of the human patella/knee region. A-B) The knees of two COL12A1 deficient patients were investigated by X-ray analysis. Frontal views reveal the localisation of the patella (P), tibia (T) and femur (F). X-ray analysis of the bent knees to visualize the facies patellaris femoris groove and the patella.

## Discussion

Here we report our findings on the function of collagen XII in the musculoskeletal system. In our analysis an overall reduction in body size and weight in the *Col12a1*^*-/-*^ mice was observed. Reduced muscle mass was detected for the gastrocnemius and quadriceps muscles. Interestingly, in the quadriceps muscle a slight increase in the number of centralized nuclei was found. However, only < 5% of the fibers have centralized nuclei and no fibrotic signs were visible in muscle sections. Further, no progressive skeletal muscle phenotype occurs, as seen in other mouse models for myopathy (17) (18). This recapitulates the phenotype observed in COL12A1 deficient patients, since the muscle phenotype is not progressive but improves during childhood (19). This indicates that collagen XII is important for muscle development and less for muscle homeostasis in adults. Expression analysis revealed that in chick embryonic muscle, collagen XII is deposited in perimysium and endomysium (3). In contrast, in adult mice collagen XII immunofluorescence staining is mainly restricted to the endomysium and scarce in perimysium. Since only few patients with COL12A1 mutations have been described, the full phenotypic spectrum of this disease is not yet defined. One clear observation is that mutations in COL12A1 often cause a strong rectus femoris atrophy (19) (6), as we have now found also in *Col12a1*^*-/-*^ mice.

An interesting hallmark of young *Col12a1*^*-/-*^ mice is the frequent dislocation of the patella. Since the facies patellaris femoris groove is not properly formed, the patella is not kept in the proper position and therefore easily slips to the side. At birth, few animals show this pathology, but upon increased physical activity during maturation up to 75% of the *Col12a1*^*-/-*^mice exhibit a patella subluxation. Furthermore, it has been described that aberrant fibril packing with altered fibril growth in *Col12a1*^*-/-*^ mice results in irregular collagen fiber cross-sectional profiles (20) and, in flexor digitorum longus tendons, in a significantly larger cross-sectional area than in wild type animals (Izu et al., 2021). The *Col12a1*^*-/-*^ flexor digitorum longus also shows an increased stiffness, a change that may contribute to the patella subluxation. Since members of the small leucine-rich proteoglycan family, e.g. decorin, lumican and fibromodulin, are also known to influence the collagen fibril morphology, it is interesting to compare our results to those from lumican and fibromodulin deficient animals. Lumican-deficient mice have collagen fibrils with increased diameter forming a disorganized matrix in cornea and skin, with a consequent decrease in corneal clarity and increased skin laxity (21). Fibromodulin deficient mice have more immature, small diameter tendon collagen fibrils (22). In comparison to *Col12a1* deficiency, double ablation of both lumican and fibromodulin genes leads to strong Ehlers-Danlos syndrome phenotype. In particular, patella subluxation and a significant reduction in tendon stiffness was observed (23). Since only one picture and no MRI or X-ray data were published by these authors, it is not clear whether the facies patellaris femoris groove is affected in the double KO mice (23). In *Col12a1*^*-/-*^ animals the tendons are stiffer and therefore the mechanism of the luxation may be different in the two mouse lines.

To study the molecular changes, single-cell RNA-seq was performed on hind limbs from 7 days old mice. Interestingly, reduced expression from different matrix genes was observed. For some regulated genes a functional connection with Ehlers-Danlos syndrome and with the development of tendon and patella can be made. For example, WIF1, a secreted protein which inhibits WNT signaling through binding to WNT, is down regulated in *Col12a1*^*-/-*^ mice. Mutations in WIF1 have been described as a potential novel cause of a Nail-Patella-like disorder (24). 95% of the cases with this disorder are attributed to mutations in the transcription factor LMX1B, causing an absent/hypoplastic patella. The LMX1B protein is important during early embryonic development of limbs, kidneys, eyes and patella. However, no changes in the facies patellaris femoris groove have been reported. Furthermore, the expression of two small leucine-rich proteoglycan, fibromodulin and keratocan, is downregulated. Keratocan is known to be important for the corneal shape (25). Additional genes regulated in *Col12a1*^*-/-*^ mice are linked to angiogenesis. Among those, CCN1, previously known as CYR61, is important for vascular morphogenesis during development, but also for tissue regeneration and fibrosis (26).

In human COL12A1-related myopathic Ehlers-Danlos syndrome both recessive and dominant modes of inheritance have been described (7). Generally, recessively inherited, bi-allelic loss-of-function variants cause a severe congenital disease characterized by hypotonia, global muscle weakness and atrophy, and joint hyperlaxity (8). In heterozygous patients, variants with a dominant-negative pathogenic effect on collagen XII fibrillar assembly have been described. These cause a much milder phenotype, characterized by mild motor developmental delay, mild proximal weakness, and joint hyperlaxity (5) (27). Ehlers-Danlos syndrome patients in general show joint dislocations and subluxations (partial dislocations) due to changes in elasticity. Additional analysis of COL12A1 patients now revealed morphological alterations of the facies patellaris femoris grooves, similar to those identified in the *Col12a1*^*-/-*^ mice. In one instance the groove/furrow was almost absent, whereas in the remaining knees the grooves seemed to be less pronounced than in unaffected individuals. However, the facies patellaris femoris grooves in COL12A1 haploinsufficiency patients should be studied further. Unfortunately, presently available patient data are not sufficient for this purpose.

In summary, our results suggest that extracellular and cellular mechanisms mediated by collagen XII are important for the correct establishment of the tendon – patella – muscle unit in the knee of mice and humans. Findings in collagen XII deficient mice lead to the identification of novel overseen phenotypes in patients. This mouse phenotype-to-patient approach was very successful, despite differences in mechanical loading of joints, to define a common phenotype between species. Further studies in mice are warranted to elucidate the precise function of collagen XII in the muscle, tendon and bone unit and to understand the pathology in patients. The patella luxations can be attributed to either muscle atrophy, changes in the tendon mechanics and/or alterations of the facies patellaris femoris groove. Most likely a combination of these factors contributes to this phenotype.

## Material and methods

### Animals

*Col12a1*^*-/-*^ mice obtained through heterozygous breedings were kept on a mixed C57BL6/N and 129/SvJ background (9). Female mice were studied at 1, 3, and 6 months of age. At P7 or less, both genders were used for experiments. All experiments were performed in agreement with the guidelines of the German animal protection law (Institutional review board:”Landesamt für Natur, Umwelt und Verbraucherschutz Nordrhein-Westfalen”; AZ: 84-02.04.2019.A326). Animals were housed in a specific pathogen-free facility at 20 to 24°C on a 12 to 12 h light/dark cycle in individual ventilated cages and supplied with standard irradiated mouse chow and water ad libitum.

### Histology and immunohistochemistry

Right hind legs of *Col12a1*^*-/-*^ mice and littermate controls were collected at the indicated time points and fixed immediately in 4% paraformaldehyde for 24 hours. Decalcification was carried out in 0.5 M EDTA (pH=8.0) at 4°C for 2 weeks with mild agitation. Samples were embedded in paraffin and sectioned by microtome (Leica Biosystem, Germany) at 7 μm. Hematoxylin-Eosin and Alcian Blue staining was performed for visualisation. Immunofluorescent staining of collagen XII was performed as follows: after deparaffinization, samples were treated with 5 mg/ml hyaluronidase (H3506, Sigma-Aldrich) in 0.1 M NaH_2_PO4, 0.1 M sodium acetate (pH=5.0) at 37°C for 30 minutes. Then samples were washed three times in TBS and incubated in 0.25% Triton X-100 (v/v in TBS) for 10 minutes at room temperature for permeabilization. After another three TBS washing steps, blocking was done at room temperature for 45 minutes with 10% FBS, 5% NGS, v/v in TBS. First antibody was diluted in 5% FBS (v/v in TBS) and incubated at 4°C overnight. The next day, sections were washed three times in TBS and incubated with secondary antibody diluted in 1% FCS/5% NGS (v/v in TBS) for 1 hour at room temperature. After another three washes in TBS, nuclear staining was performed by applying DAPI (4 mg/mL in TBS, 32670, Sigma) at room temperature for 5 minutes.

Gastrocnemius and quadriceps muscles were dissected from 6 months old WT and KO mice and immediately snap-frozen in cold isopentane. Samples were stored at -80°C until sectioning. Muscle samples were sectioned with a cryostat (Leica, Germany) at a thickness of 7 μm and captured with Superfrost Plus adhesive microscopy slides (Th. Geyer, Germany). Hematoxylin-Eosin staining was performed according to standard methods. For the immunostaining, the slides were fixed for 10 minutes with 4% paraformaldehyde in PBS, permeabilized for 10 minutes with 0.3% Tween20 (v/v in PBS) and stained with rabbit polyclonal antibodies against collagen XII (KR144;1:200), collagen VI (1:1000, gift from Raimund Wagener, Cologne) and Alexa 488 conjugated wheat germ agglutinin (WGA) (Invitrogen, 1:500), followed by corresponding secondary antibodies coupled to Cy3 (1:800; Jackson ImmunoResearch) and DAPI (4 mg/mL in PBS, 32670, Sigma). All samples were embedded in Mowiol mounting medium and analysed by fluorescence microscopy (Nikon Europe Eclipse TE2000-U microscope).

### Statistical evaluation

Whole muscle sections stained with DAPI and WGA were stitched together with the Grid/Collection stitching tool (Preibisch et al., 2009) in the Fiji/ImageJ software (National Institutes of Health). The myosoft extension (Encarnacion-Rivera et al., 2020) for the Fiji/ImageJ was used to calculate the number of muscle fibres and the minimal Feret’s diameter. The number of central nuclei was counted using the cell counter tool in Fiji/ImageJ. The variance coefficient of the minimal Feret’s diameter was calculated (Dubach-Powell et al., 2008)

### μCT analysis

For I_2_KI-enhanced μCT scanning, a protocol (Charles *et al*., 2016) was adapted as follows: right hind legs of P7 mice were removed and frozen in -80°C immediately after sacrifice. The limbs were then thawed and placed in 10% neutral-buffered formalin (NBF) (HT501128-4L, Sigma) for 24 hours at 4°C under constant rotation for soft tissue fixation. Then samples were washed in PBS to remove residual fixative. Samples were first scanned in a SkyScan 1172 system (Bruker microCT, Belgium) with source settings of at 50 kV, 200 μA, and a 960 ms exposure time and a pixel size of 4.98 μm using an Aluminium 0.5 mm filter. After CT scanning, the specimens were immersed in iodine-potassium iodide solution (I_2_KI, Lugol’s solution, Sigma, L6146) for eight days to enhance soft tissue contrast, and placed in 70% ethanol solution until μCT scanning using the SkyScan 1172 system at the following settings: 60 kV, 167 μA, 4.98 μm resolution, Aluminium 0.5 mm filter and 960 ms exposure time. The images were reconstructed using NRecon software (Bruker microCT, Belgium), whereas volume renderings were constructed using CTvox software (Bruker microCT, Belgium). For μCT scanning the knee joints of the right hind legs from 1 month old female mice were isolated, immediately fixed in 70% EtOH and stored in 4°C. Knee joints were evaluated using a high-resolution μCT scanner (μCT 35; Scanco Medical AG) and were scanned with an isotropic voxel size of 6 × 6 × 6 μm using 70-kVp tube voltage, 114-μA tube current, and 400 ms integration time. Image noise was removed by preprocessing of the grayscale data of the raw CT images using a 3D Gaussian filter algorithm and separation of mineralized tissue from the soft tissue by a global threshold of 200.

### Single-cell RNA-seq

Left hind legs of P7 mice were isolated from one WT and one *Col12a1*^*-/-*^ mouse and digested in 4 mg/mL collagenase II (Worthington, USA) in DMEM/F-12 (supplied with 10% FBS) for 6 hours. After digestion, isolated cells were passed through 100 μm and 40 μm cell strainers (BD Bioscience, USA) to obtain single cell suspensions. Cells were washed three times with 0.04% bovine serum albumin (Serva Electrophoresis GmbH, Germany) (v/v in PBS) by centrifugation at 400 g for 5 min and brought to a density of 1,000 cells/μL. Cell viability (>90%) was confirmed by 0.4% trypan blue staining prior to processing according to 10x Genomics Chromium Single Cell 3’ Reagent Kit User Guide (v3.1). Briefly, approximately 5,000 cells of each genotype were loaded into Chromium Next gel beads in emulsion (GEMs) Chip G and single cell capture was performed with 10X Chromium Controller (10x Genomics). cDNA generation and library preparation were performed using Chromium Single Cell 3’v3.1 reagent kit. Cell Ranger software (v3.1.0, 10X Genomics) were used to align reads, generate feature-barcode matrices, perform clustering and the data imported and analyzed by Loupe Browser (v5.0, 10X Genomics).

### Matrisome analysis

Matrisome gene clusters, annotated in the matrisome database 2.0 (http://matrisomeproject.mit.edu) were imported and selected for hierarchical cluster analysis to determine relationships among the expression levels using Instant clue software (Nolte et al, 2018). A fold change cut off (FC ≥ 1) and false discovery rate cut off (p-value ≤ 0.1) was used to define differentially expressed mRNAs between selected groups. False discovery rate was adjusted using the Benjamini-Hochberg procedure. Expression intensity plots (MvA plots) were generated by highlighting regulated matrisome genes within the entire entities. The entity lists were exported to generate graphical representations as Venn diagrams using the FunRich 3.1 tool (28).

## Acknowledgements

We would like to thank the patient families for their participation and support of this research. In addition, we thank Semra Oezcelik and Christian Frie for excellent technical assistance.

