## Supplement for "Ablation of the FACIT collagen XII disturbs musculoskeletal ECM organization and causes patella dislocation and myopathy"

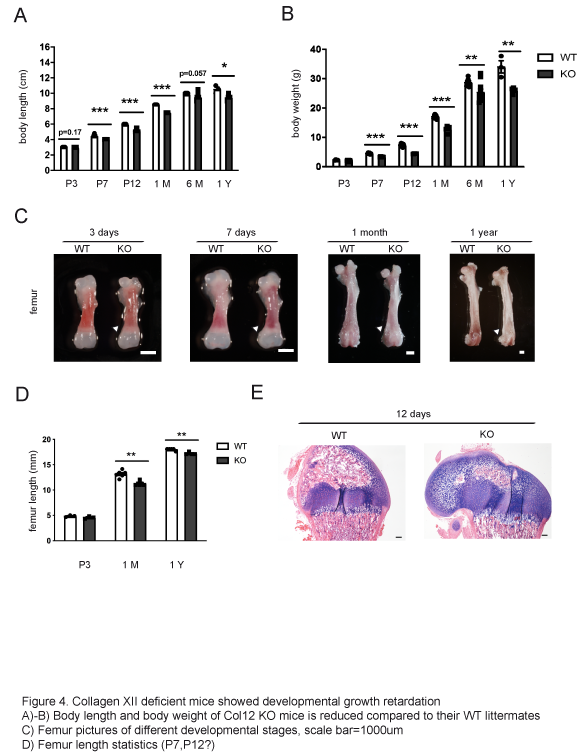

**Suppl. Fig 1. *Col12a1^-/-^* mice show growth retardation.** *Col12a1^-/-^* mice show a reduced body length (A) and weight (B) compared to their WT littermates. A shorter femur was found in *Col12a1^-/-^* mice (C,D), scale bar 1 mm. (E) H&E and Alcian Blue staining shows that the secondary ossification center formation is delayed at P12 in *Col12a1^-/-^* mice. Scale bar: 100 µm.

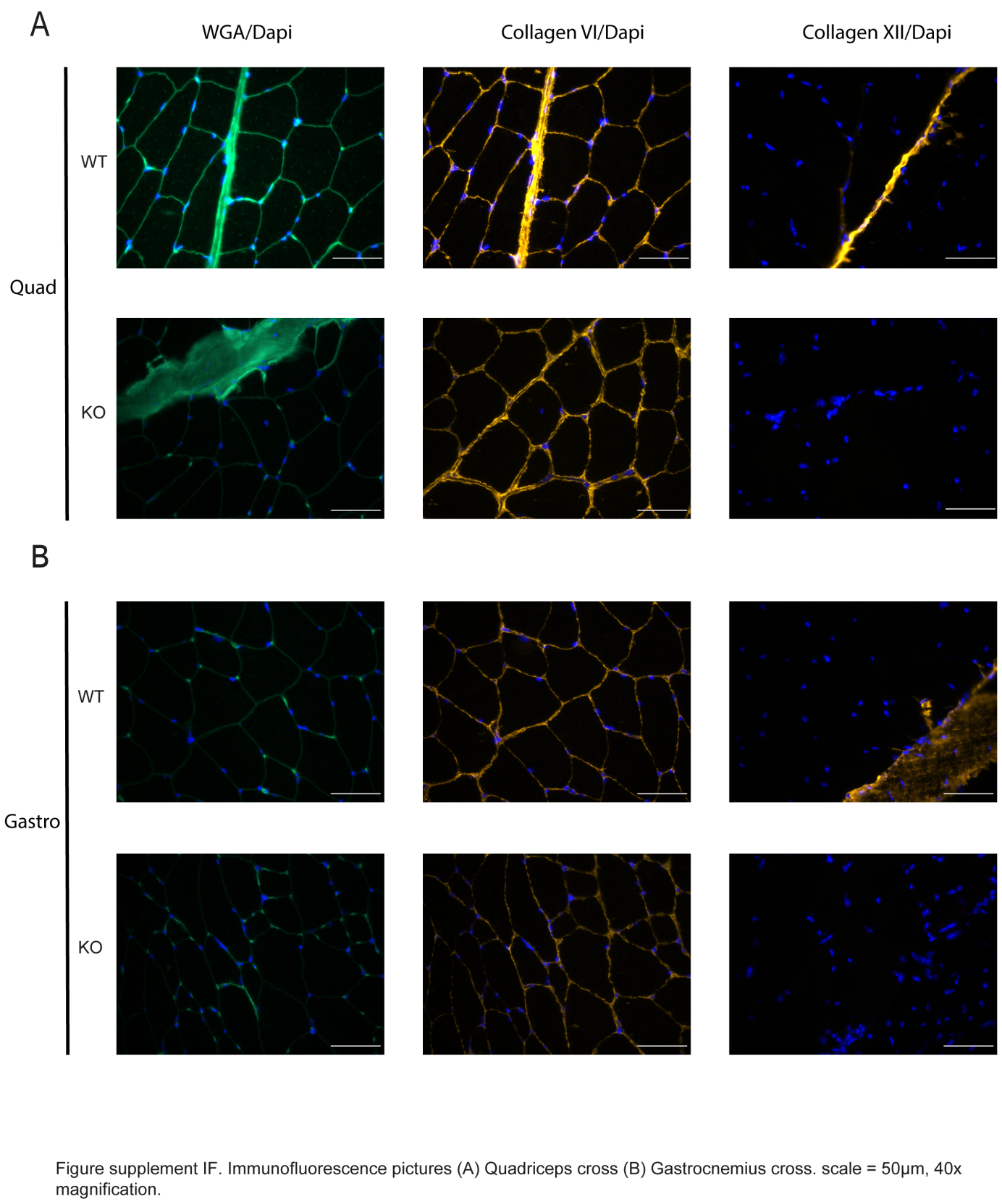

**Suppl. Fig 2. Immunofluorescent staining of the quadriceps and gastrocenemius muscles.** Fluorescently tagged wheat germ agglutinin (WGA) lectin visualizes the connective tissue in muscles. Collagen VI and collagen XII was stained in both the quadriceps and gastrocnemius muscle group of 6 months old mice of both genotypes (n=4).

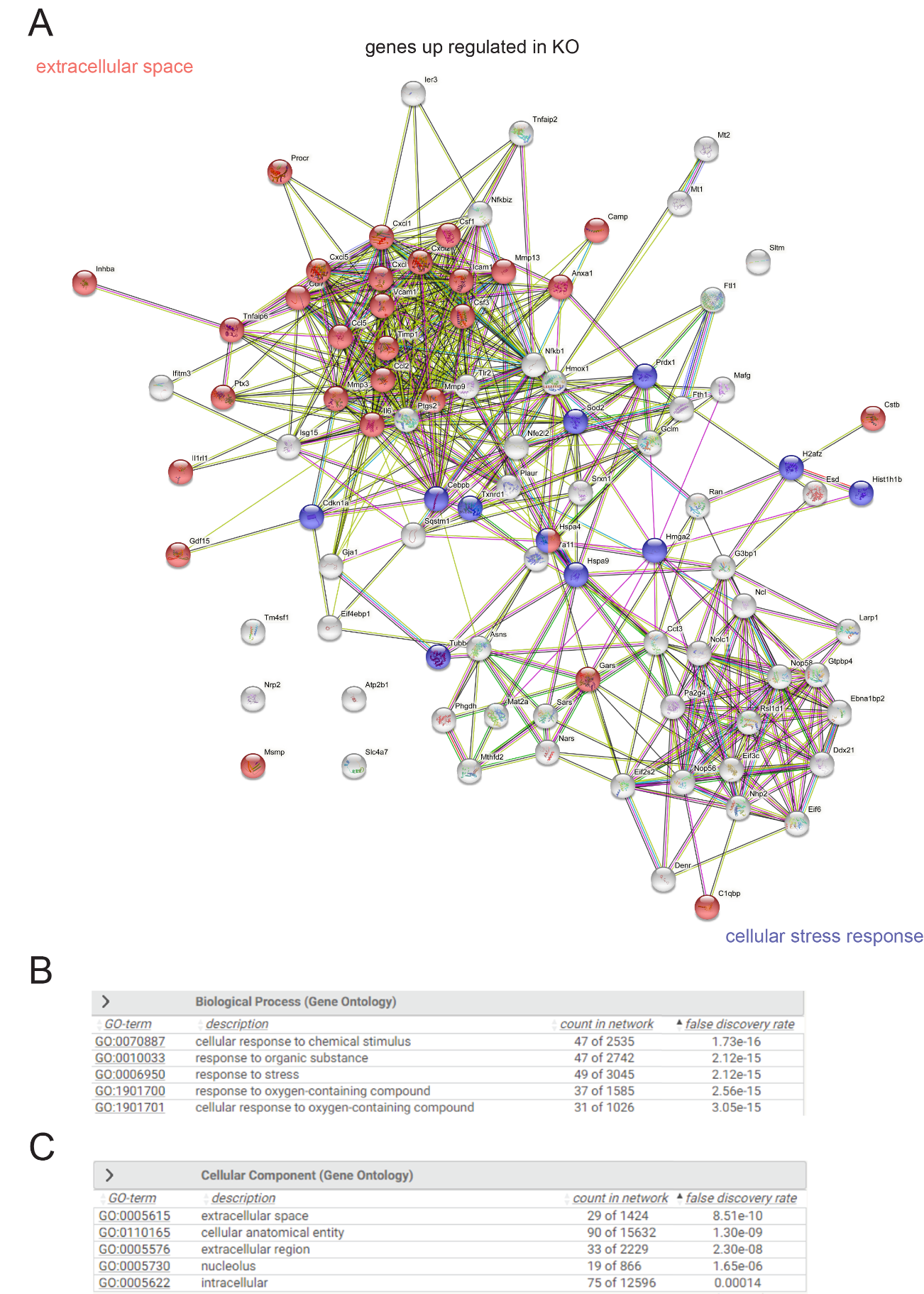

**Suppl. Fig 3. String analysis of up regulated genes in *Tnmd+* cell population from a *Col12a1^-/-^* mouse*.*** All 388 genes from the meta‐signature were used as input for STRING analysis and a network was built based on high confidence (0.8) evidence from experimental protein‐protein interaction (blue lines) and curated (purple lines) databases. Proteins are indicated by nodes labeled with the encoding gene symbol. Two genes (CEBPA and SEPP1) are not present in the used STRING version 9.1. The network is enriched in interactions (p=0.007) using the intersection of 8,612 genes present on all platforms as background.

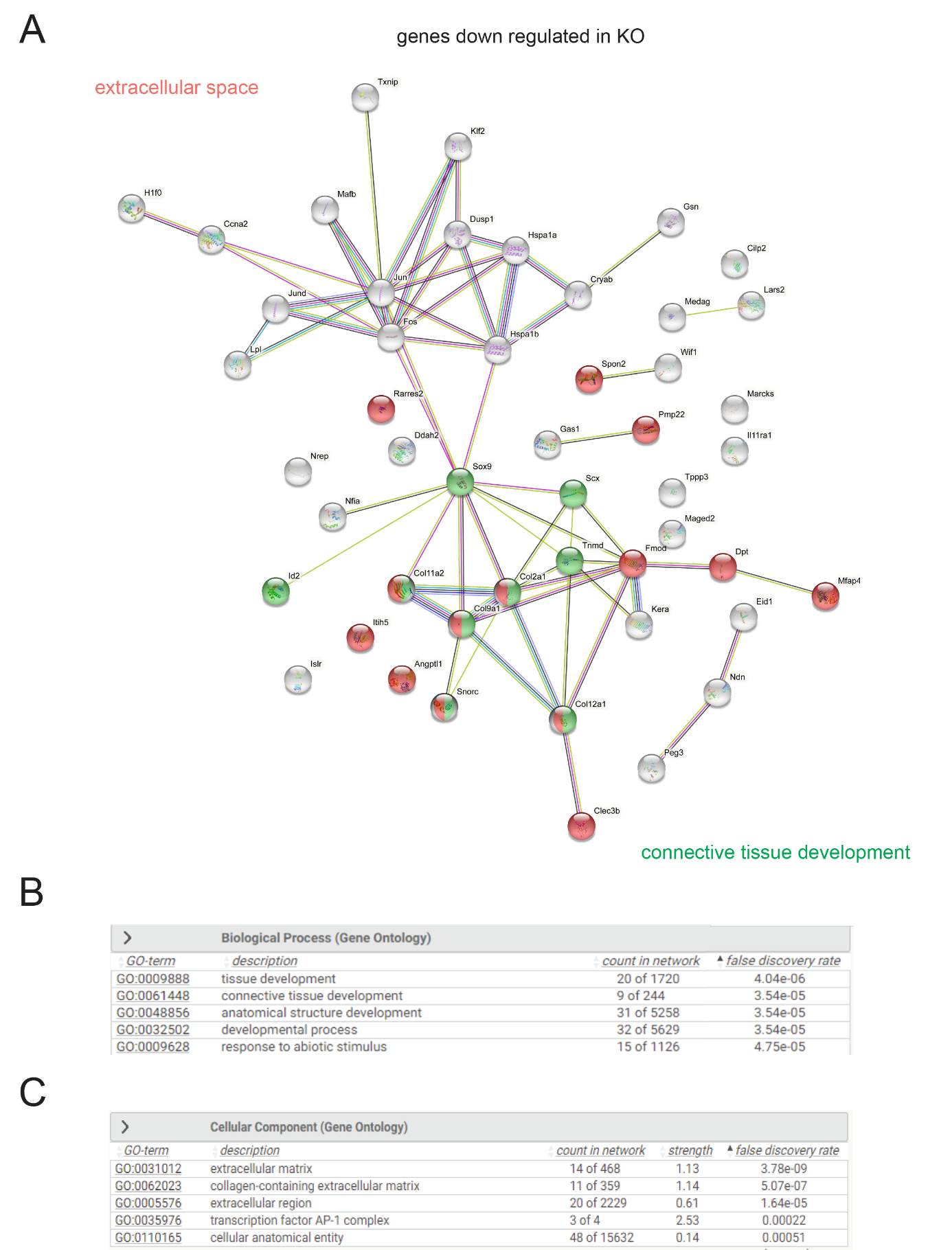

**Suppl. Fig 4. String analysis of down regulated genes in *Tnmd+* cell population from a *Col12a1^-/-^* mouse*.*** The STRING analysis and a network was built based on high confidence (0.8) evidence from experimental protein‐protein interaction (blue lines) and curated (purple lines) databases. Proteins are indicated by nodes labeled with the encoding gene symbol.

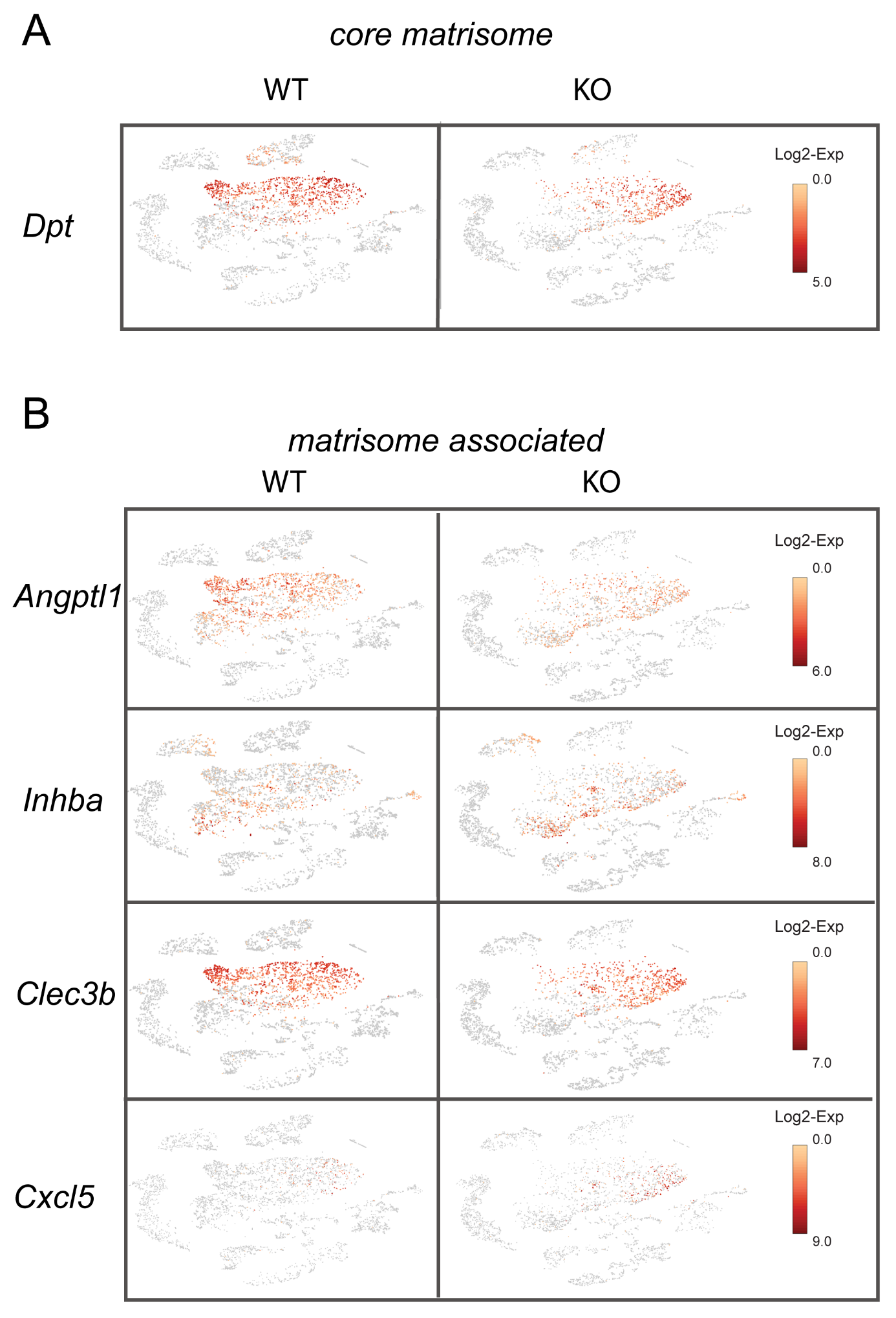

**Suppl. Fig 5. Additional candidates selected in the matrisome analysis of the *Tnmd+* cell subpopulation.** A) The proportions of entities within the core matrisome or matrisome associated cluster (B) are shown in a Venn diagram.
